## Supplementary figures and images for "Distinct Hierarchical Alterations of Intrinsic Neural Timescales Account for Different Manifestations of Psychosis"

### Supplemental Data 1

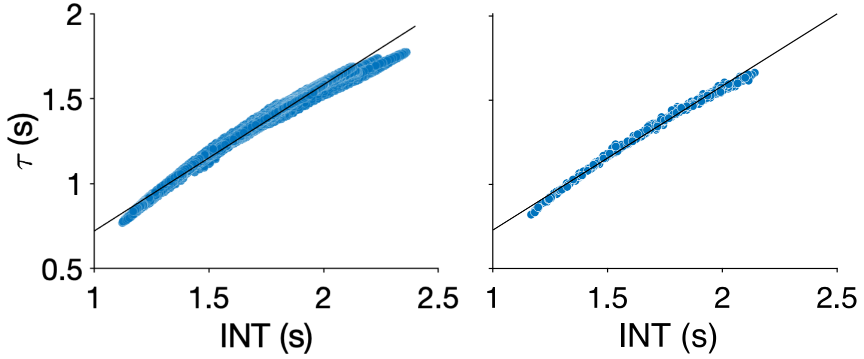

### Supplemental Data 2

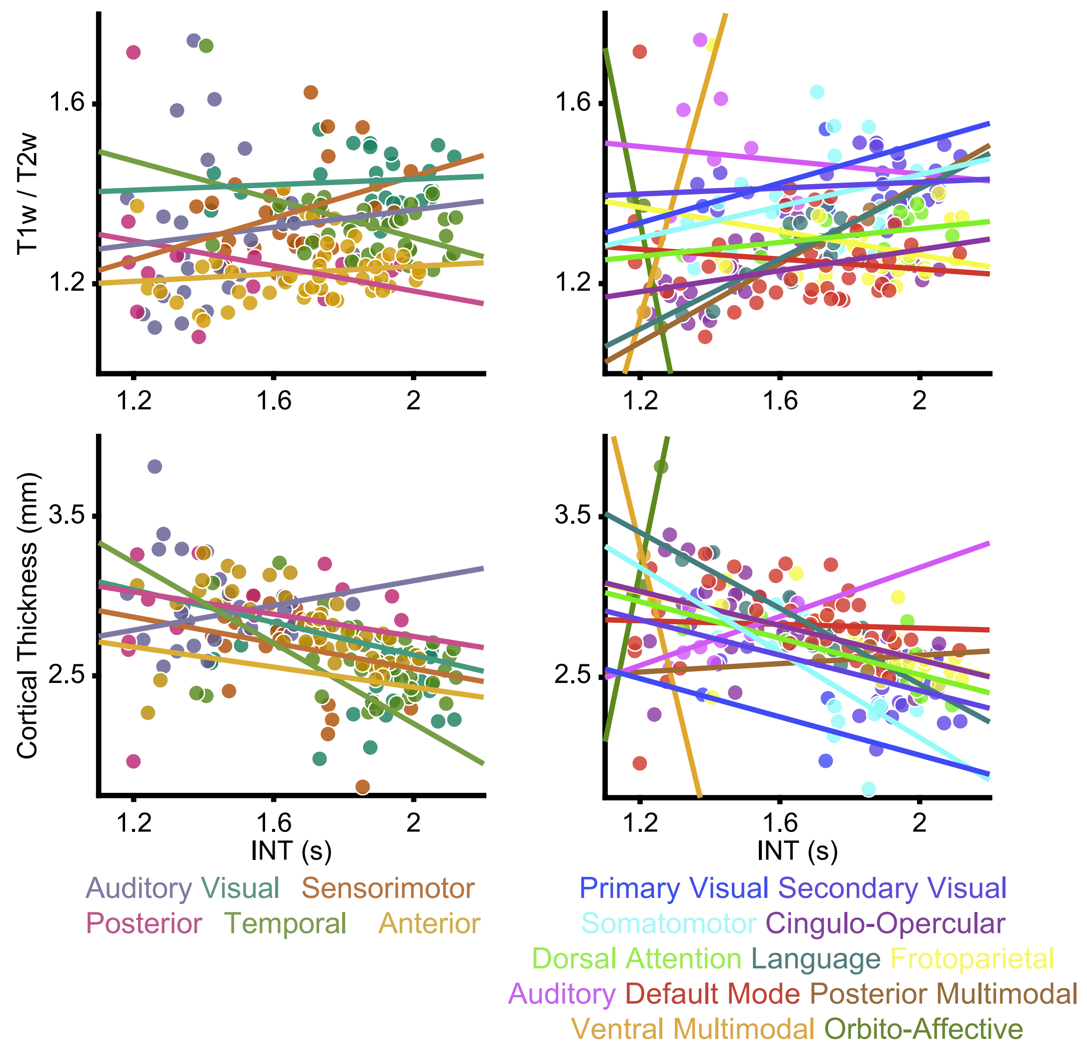

### Supplemental Data 3

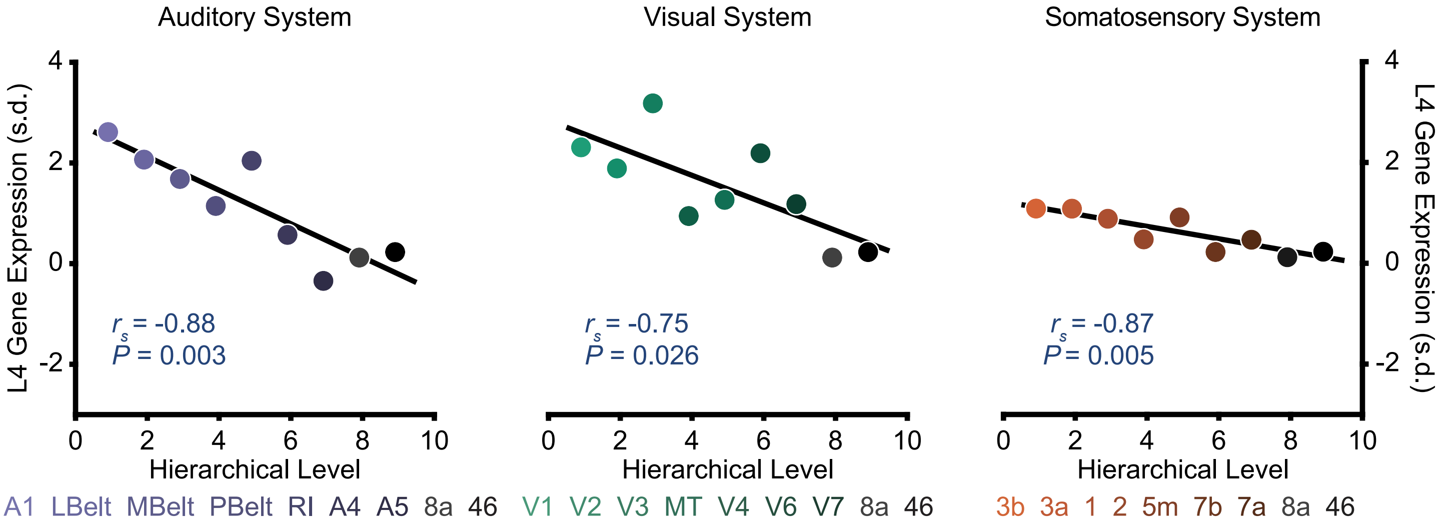

### Supplemental Data 4

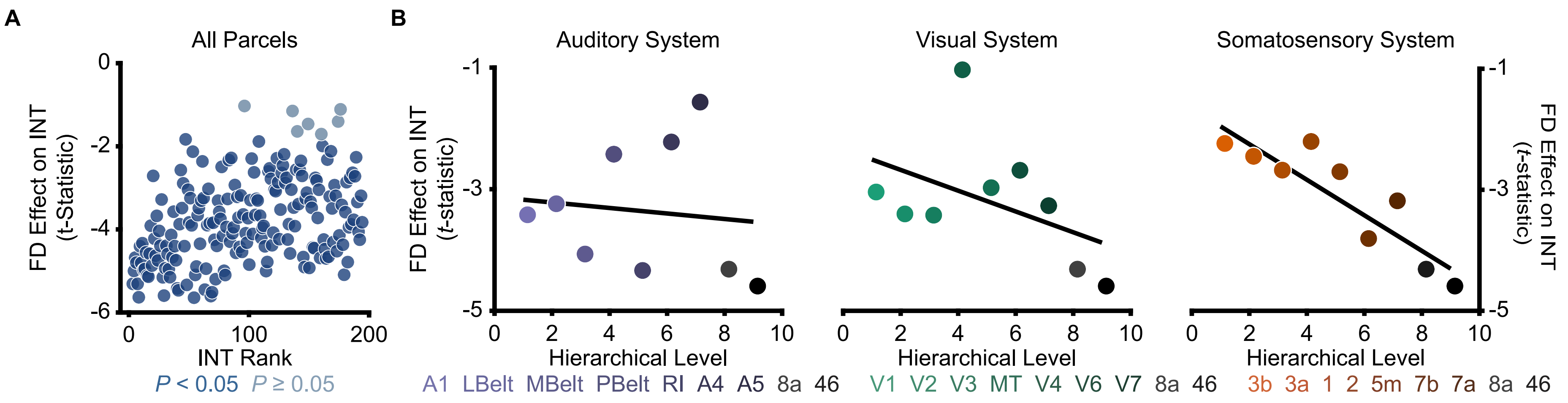

### Supplemental Data 5

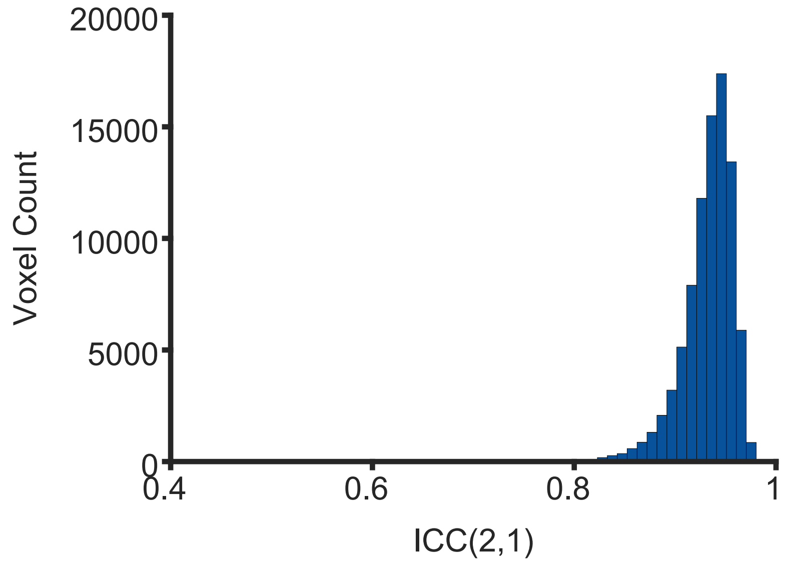

### Supplemental Data 6

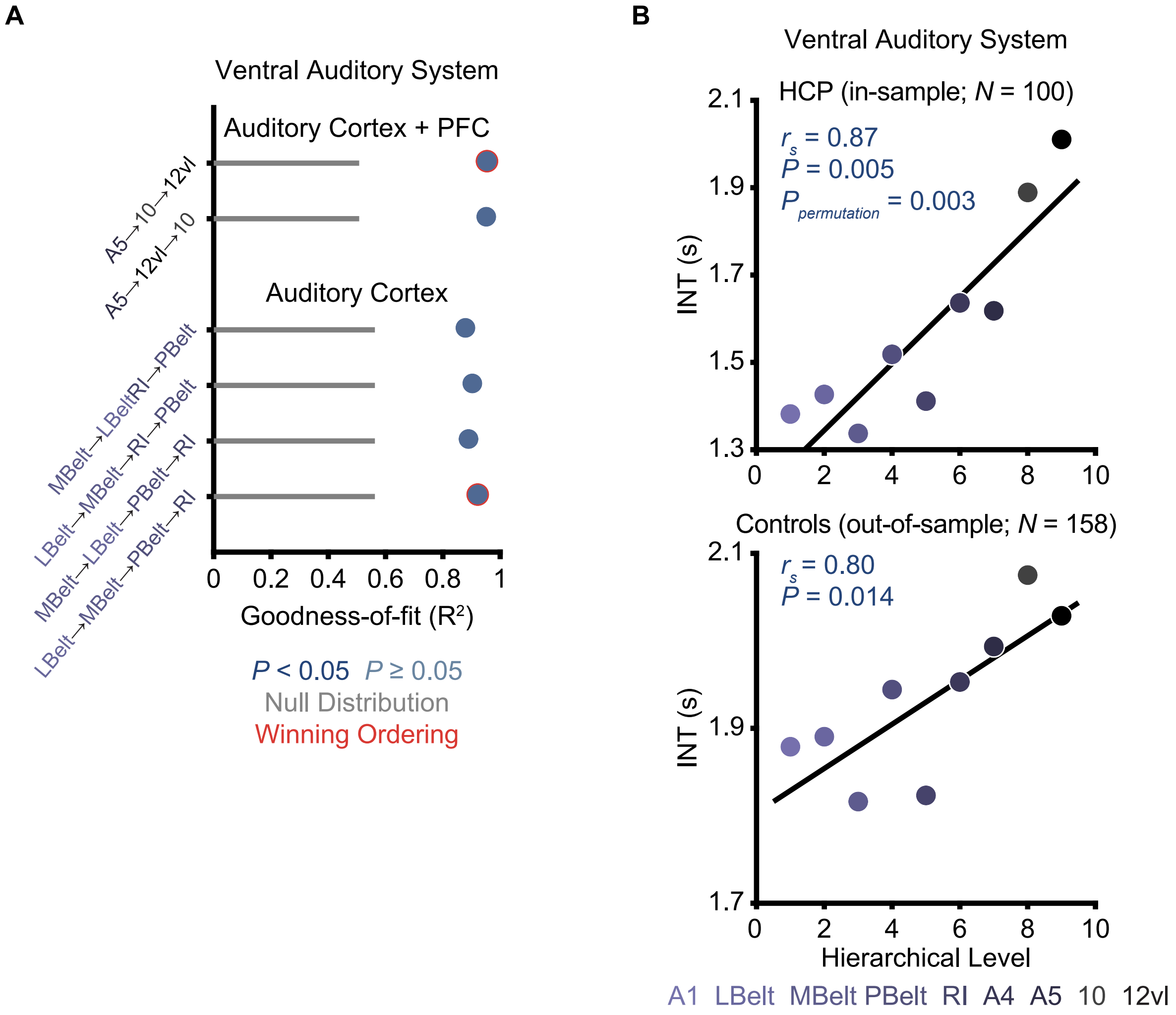

### Supplemental Data 7

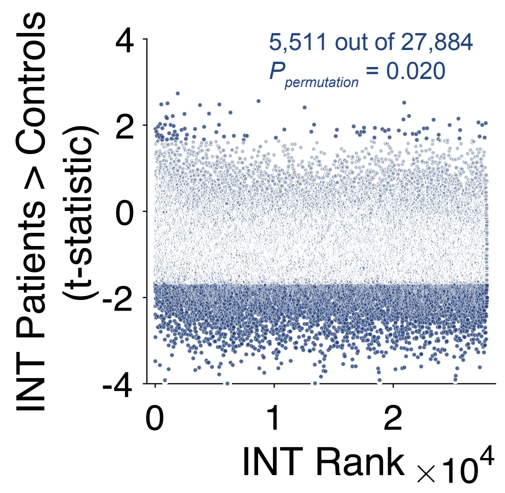

### Supplemental Data 8

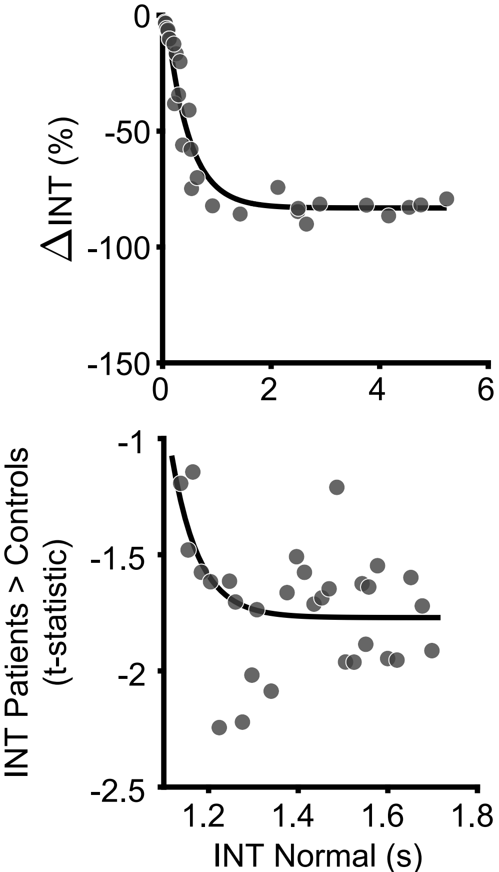

### Supplemental Data 9

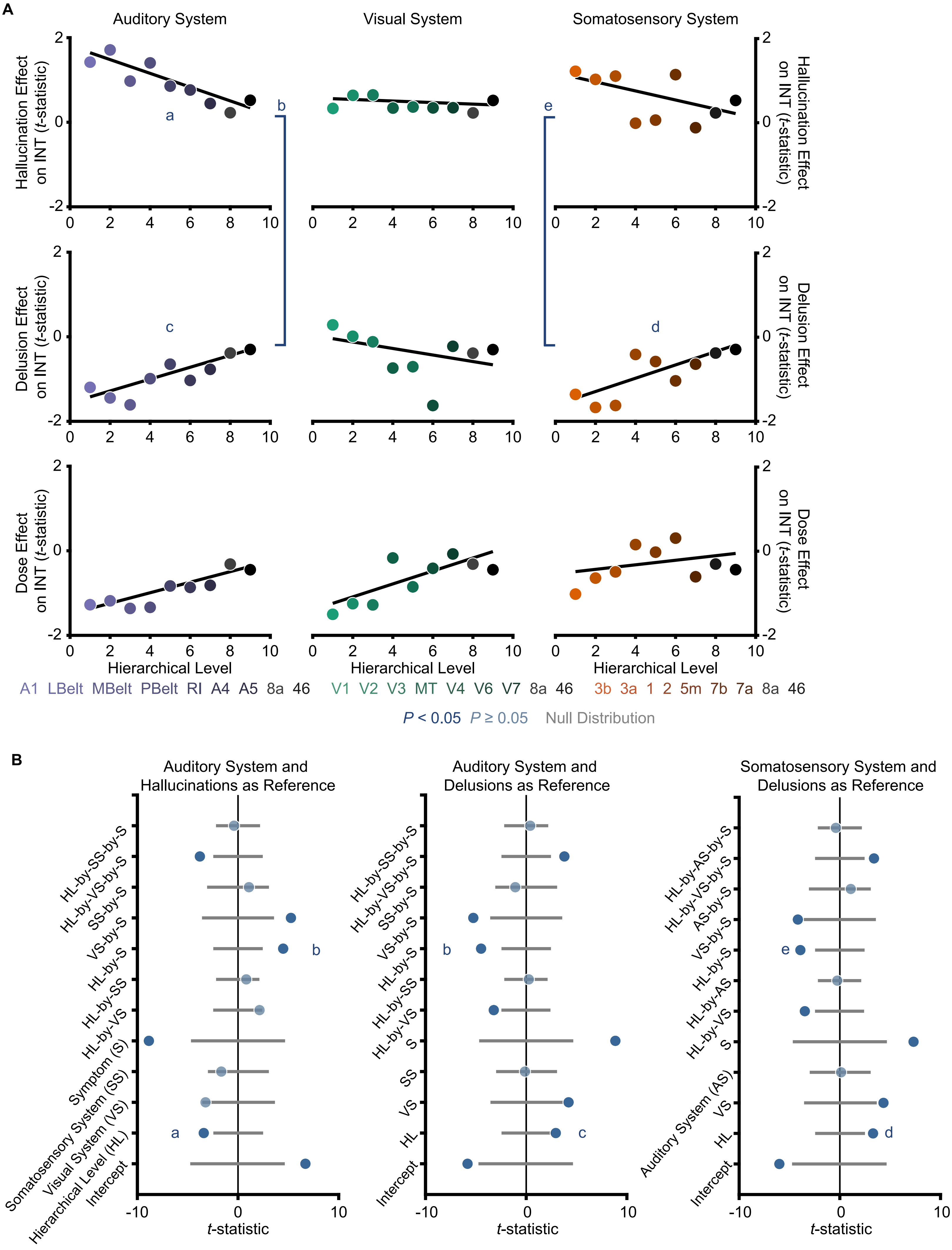

### Supplemental Data 10

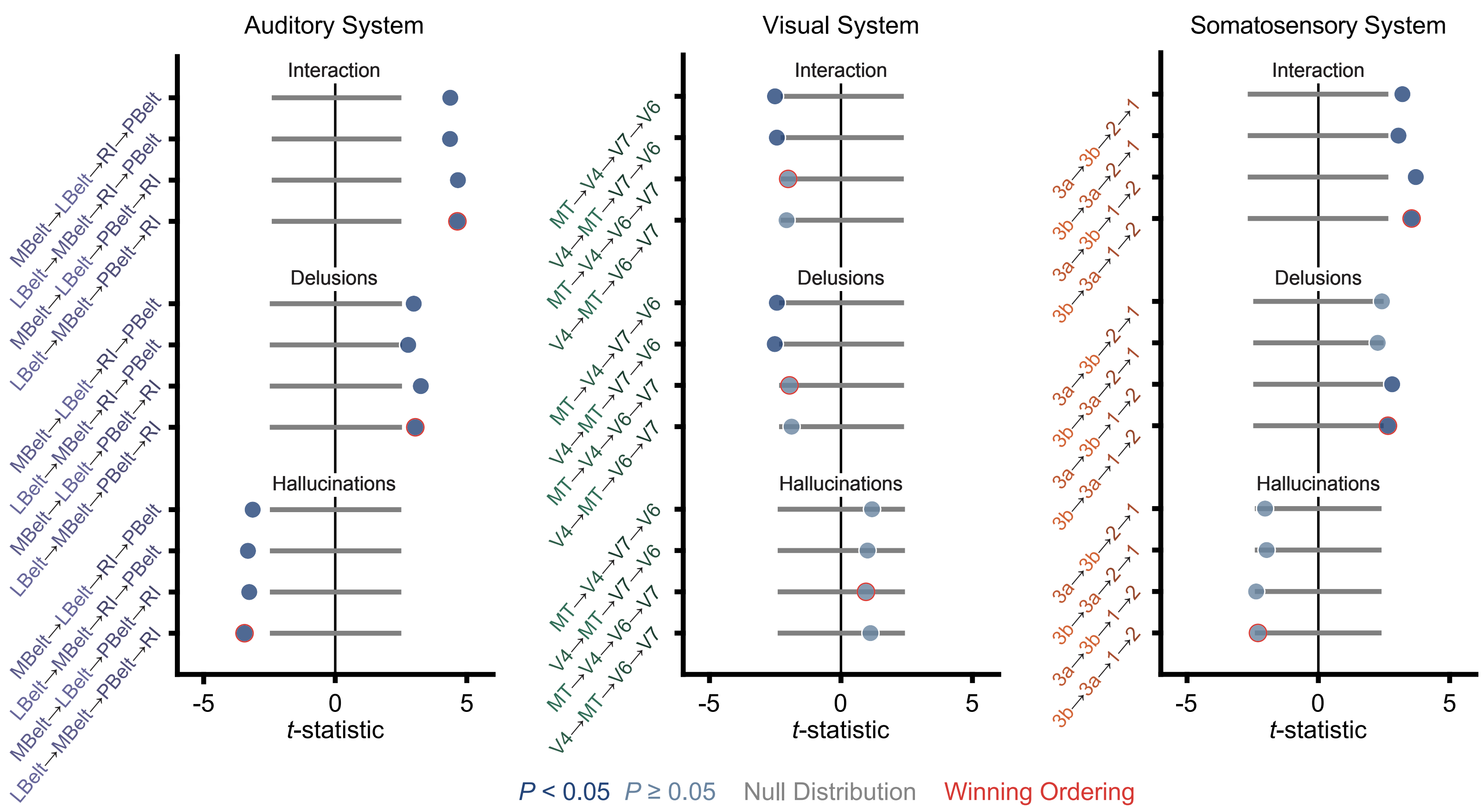

### Supplemental Data 11

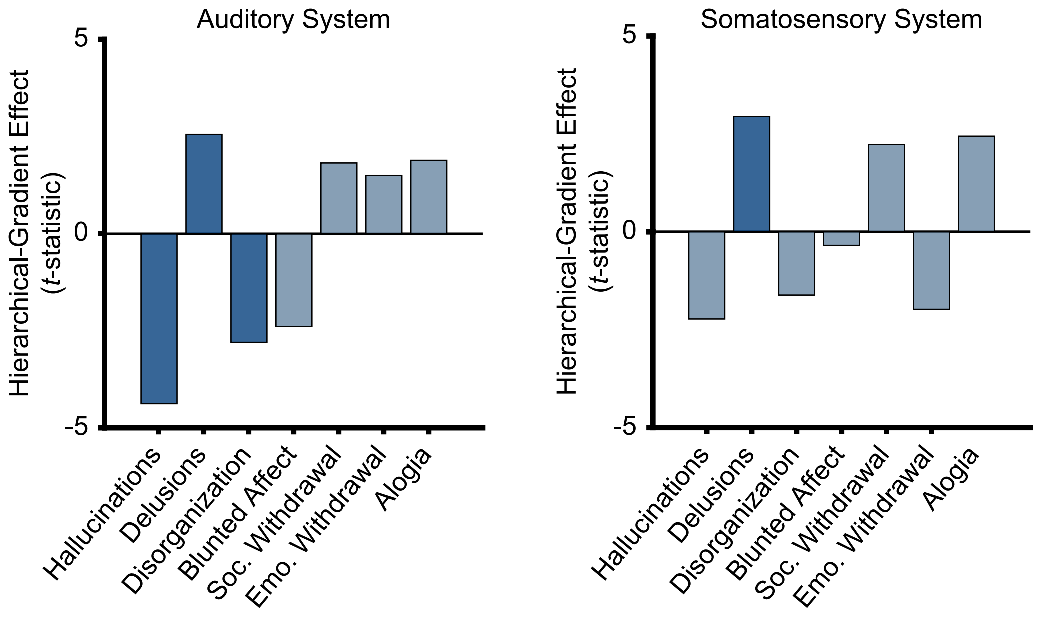

### Supplemental Data 12

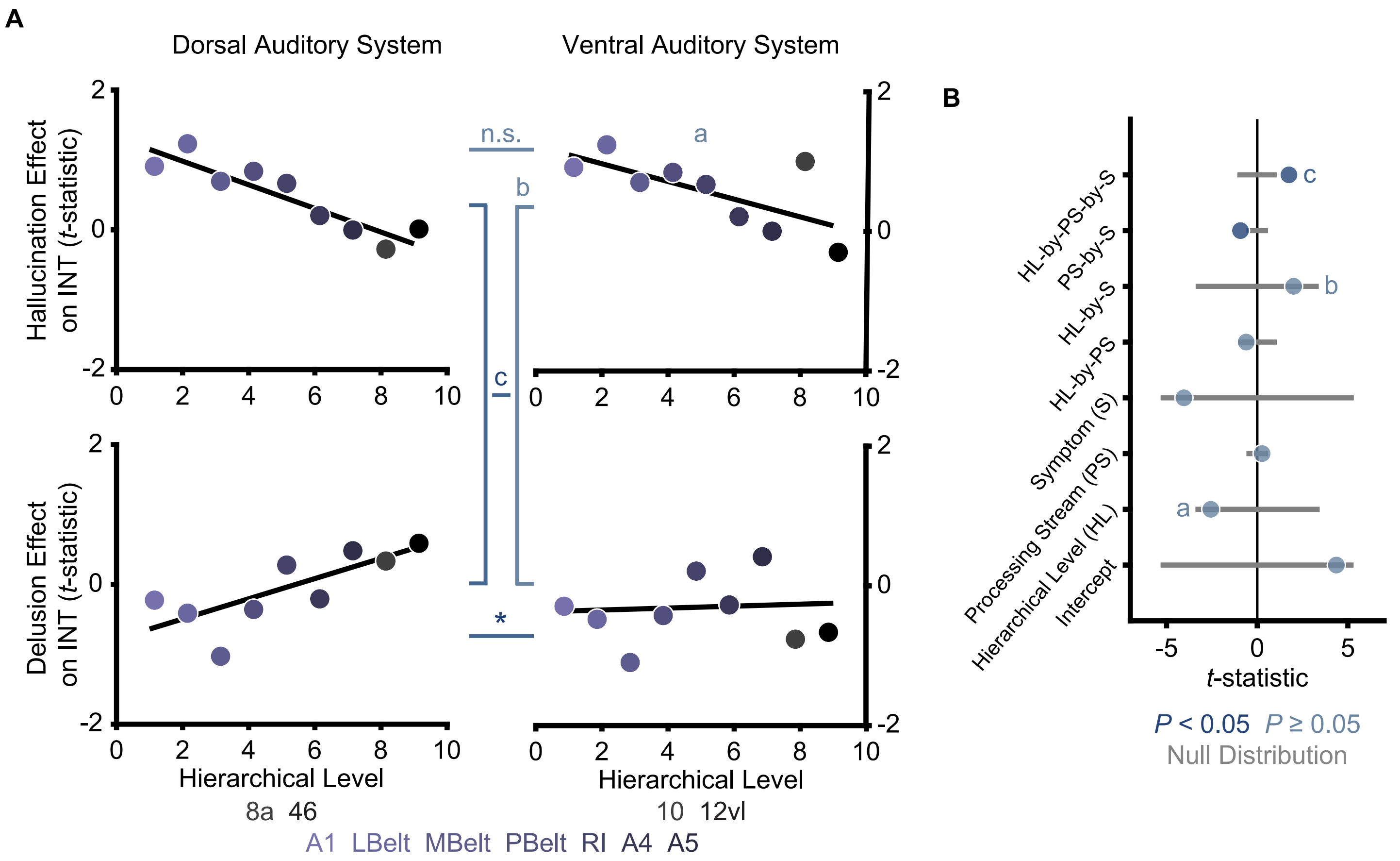

### Supplemental Data 13

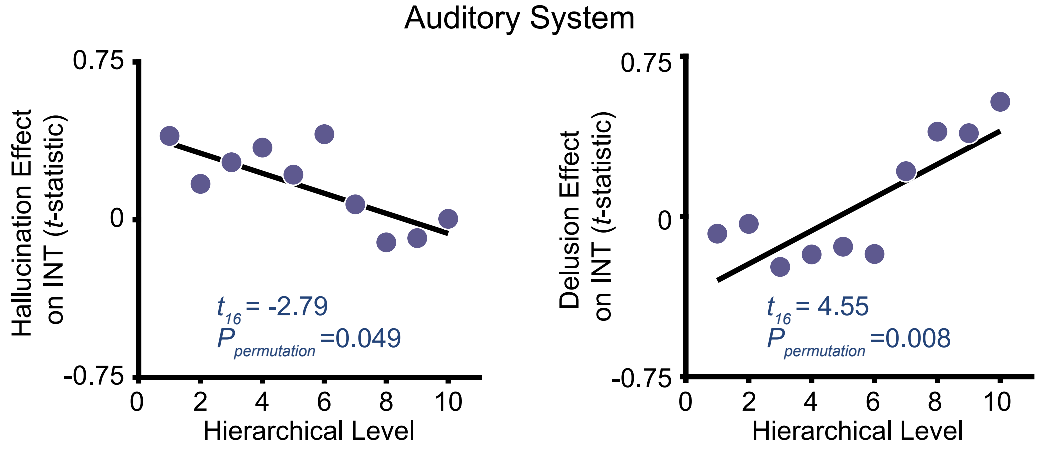
